# Differential Nr4a1 and Nr4a3 expression discriminates tonic from activated TCR signalling events in vivo

**DOI:** 10.1101/767566

**Authors:** Emma Jennings, Thomas A.E. Elliot, Natasha Thawait, Shivani Kanabar, Juan Carlos Yam-Puc, Masahiro Ono, Kai-Michael Toellner, David C. Wraith, Graham Anderson, David Bending

## Abstract

*Nr4a* receptors are activated by T cell receptor (TCR) and B cell receptor (BCR) signalling and play key roles in T cell differentiation and promoting T cell exhaustion. How TCR signalling pathways regulate Nr4a receptors and their sensitivities to different physiological types of TCR signalling (e.g. tonic versus activating) remains unknown. Here we utilise Nr4a1/Nur77-GFP and *Nr4a3*-Tocky mice to elucidate the signalling pathways that govern Nr4a receptor expression in CD4^+^ and CD8^+^ T cells. Our findings reveal that *Nr4a1-3* are Src family kinase-dependent. Moreover, *Nr4a2* and *Nr4a3* are abolished by calcineurin inhibitors and bind NFAT1, highlighting a necessary and sufficient role for NFAT in the control of *Nr4a2* and *Nr4a3*, but redundancy for NFAT for *Nr4a1*. During T cell development, Nr4a1 is activated by tonic signalling during TCR-beta selection in the thymus, whilst Nr4a3 requires cognate peptide:MHC interactions for expression. Thus, due to differential sensitivity of Nr4a1 and Nr4a3 to TCR signalling pathways, T cells undergoing tonic versus activating TCR signalling events can be distinguished in vivo.

## Introduction

The Nr4a family of orphan nuclear receptors consists of three members: Nr4a1 (Nur77), Nr4a2 (Nurr1) and Nr4a3 (Nor1). Nr4a receptors have been shown to be ligand-independent and their structure is set to a constitutively active form (Wang et al., 2003). Nr4a1 and Nr4a3 have been shown to be rapidly upregulated in peripheral T cells (Ashouri and Weiss, 2017; Bending et al., 2018b; Moran et al., 2011; Zikherman et al., 2012) and thymocytes (Cheng et al., 1997), following T cell receptor (TCR) signalling, and are considered a more specific marker of T cell activation than the cell surface marker CD69, which can be upregulated by non TCR stimuli (Moran et al., 2011). Expression of Nr4a1 (Moran et al., 2011) and Nr4a3 (Bending et al., 2018b) in CD4^+^ T cells is lost in major histocompatibility complex (MHC) Class II knockout mice in vivo, highlighting the key role of TCR signalling in regulating expression of the Nr4a receptors.

Early studies of Nr4a1 identified a key role for it in the induction of apoptosis by TCR-induced Nur77 protein expression (Liu et al., 1994), which associates with Bcl-2 at the mitochondria to mediate negative selection (Thompson and Winoto, 2008). Nr4a2 can bind *Foxp3* regulatory elements and regulate the differentiation of CD4^+^ T cells (Sekiya et al., 2011), and persistent *Nr4a3* expression is a hallmark of Treg undergoing differentiation in the thymus (Bending et al., 2018b). Genetic ablation of Nr4a receptors abolishes Treg development and can lead to multiorgan autoimmunity (Sekiya et al., 2013). Nuclear Nr4a1 can modulate Treg differentiation and clonal deletion (Fassett et al., 2012), highlighting a critical role for Nr4a family members in thymic T cell development and immune regulation.

More recent studies of the Nr4a family have found that nuclear factor of activated T cells (NFAT) (Martinez et al., 2015) and Nr4a receptors (Mognol et al., 2017; Scott-Browne et al., 2016) are intimately linked to the development of CD8^+^ T cell exhaustion, suggesting that NFAT and Nr4a family members may cooperate during chronic antigen stimulation to adapt T cell programmes. Indeed, Nr4a family members have been shown to limit chimeric antigen receptor (CAR) T cell function in solid tumours (Chen et al., 2019) and that Nr4a1 may be a key mediator of T cell dysfunction through binding and repressing expression of the activator protein one (AP-1) transcription factor (Liu et al., 2019). Nr4a family member function is complex since they can also promote CD8^+^ T cell exhaustion through cooperation with other transcription factors such as thymocyte selection-associated high mobility group box protein (TOX) and TOX2 (Seo et al., 2019). In addition, Nr4a1 can alter the metabolic state of T cells, acting as a break during T cell activation to dampen inflammation (Liebmann et al., 2018).

Given the central roles of Nr4a members in autoimmunity and cancer, they are an emerging therapeutic target. For example, pharmacological inhibition of Nr4a nuclear receptors can enhance antitumor immunity (Hibino et al., 2018). Therefore, understanding the regulation of the Nr4a transcription factors is of both fundamental and therapeutic interest. Whilst Nr4a members are paralogues, the signalling pathways that regulate their expression have not been clearly defined, although a role for NFAT1 has been proposed for CD8^+^ T cells (Martinez et al., 2015; Mognol et al., 2017; Scott-Browne et al., 2016). Here we utilised the state-of-the-art Nr4a3-Timer of cell kinetics and activity (Tocky) system (Bending et al., 2018a; Bending et al., 2018b) and Nr4a1/Nur77-GFP Mice (Moran et al., 2011) to determine the pathways regulating *Nr4a* transcription in CD4^+^ and CD8^+^ T cells. Our findings highlight that the calcineurin (CaN)/NFAT pathway is necessary and sufficient for the induction of Nr4a2 and Nr4a3, but redundant for Nr4a1. Moreover, while Nr4a1 is activated in response to tonic T and B cell receptor signalling during lymphocyte development in both bone marrow and thymus, Nr4a3 activation in mature T- and B-cells requires a higher threshold of signalling that is achieved only through activating TCR and BCR signalling.

## Results

### Peptide stimulation induces Src family kinase and calcineurin-dependent Nr4a3 transcription in CD4^+^ and CD8^+^ T cells

In order to interrogate the signalling pathways that control *Nr4a3* expression, *Nr4a3-* Tocky mice were crossed with the Tg4 Tiger TCR transgenic line (Burton et al., 2014), which express an autoreactive CD4^+^ TCR specific for myelin basic protein (MBP) (Liu et al., 1995). This system allows a sophisticated dissection of the temporal patterns of TCR signalling in response to self-agonist peptides (Supplementary Fig. 1A). In addition, in order to study CD8^+^ TCR signalling dynamics Nr4a3-Tocky mice were bred to the ova-peptide specific OTI TCR transgenic mice (Hogquist et al., 1994) (Supplementary Fig. 1B). Analysis of spleen and thymus of these two transgenic lines revealed selective expression of the transgenic beta chains Vβ8 TCR (Tg4) and Vβ5 in OTI (Supplementary Fig. 1). Stimulation with myelin basic protein (MBP) Ac1-9[4Y] peptide (Tg4) or ova peptide (OTI) in vitro triggered activation of Nr4a3-Timer expression, resulting in early Blue fluorescence, before time dependent maturation to a Blue^+^Red^+^ fluorescent state indicative of persistent dynamics of Nr4a3 expression (Fig. 1A&B). Expression of Nr4a3-Blue (i.e. active TCR signalling) was shown to be dependent on the Src family kinase, since incubation of Tg4 Nr4a3-Tocky T cells with the inhibitor PP2 (Hanke et al., 1996) abolished peptide-induced expression of Nr4a3 and CD69 in both Tg4 (Fig. 1C) and OTI (Fig. 1D) T cells. TCR signalling results in the activation of many signalling intermediaries but converge on the activation of at least three key transcription factors: NFAT, AP1 and NFκB (Brownlie and Zamoyska, 2013). Given that membrane depolarisation and calcium flux are central to T cell activation and that Nr4a receptors have previously been linked to NFAT activation (Scott-Browne et al., 2016), we investigated the sensitivity of *Nr4a3* to NFAT pathway inhibitors. To do this we utilised two distinct inhibitors of calcineurin, a key enzyme in TCR signalling which dephosphorylates NFAT in response to Ca^2+^ signalling (Hogan et al., 2003). Interestingly, incubation with either cyclosporin A (a fungal derived product that forms a complex with cyclophilin to block the phosphatase activity of calcineurin (Matsuda and Koyasu, 2000)) or FK506 (Tacrolimus, a macrolide calcineurin inhibitor (Thomson et al., 1995)) completely abolished TCR stimulation-induced Nr4a3-Timer Blue expression in CD4^+^ Tg4 T cells (Fig. 1E&F), whilst not significantly affecting the frequency of CD69^+^ T cells (Fig. 1G). Cyclosporin A, however, significantly reduced CD69 expression levels (Fig. 1H), suggesting that CD69 is in part regulated by the CaN/NFAT pathway. Identical findings were mirrored for CD8^+^ T cells (Fig. 1I-L).

**Figure 1:**
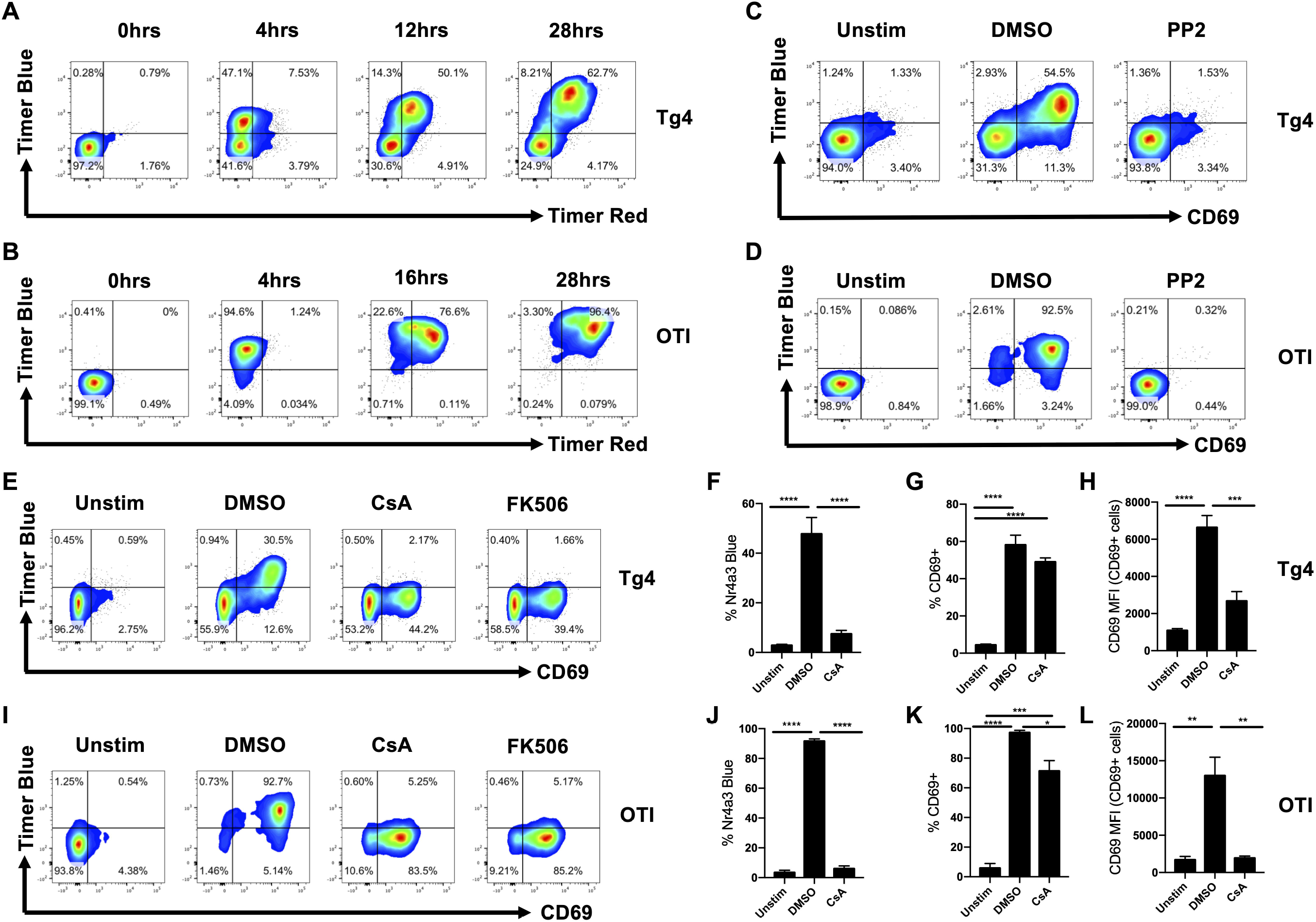
Peptide stimulation induces Src family kinase and calcineurin-dependent *Nr4a3* transcription in CD4^+^ and CD8^+^ T cells. **(A)** Splenocytes from Tg4 Nr4a3-Tocky mice were stimulated with 10μM 4Y-MBP variant for the indicated number of hours before analysis by flow cytometry for expression of Nr4a3-Timer Blue vs. Nr4a3-Timer Red in CD4^+^ Tg4 T cells. **(B)** Splenocytes from OTI Nr4a3-Tocky mice were stimulated with ova peptide for the indicated number of hours before analysis by flow cytometry for expression of Nr4a3-Timer Blue vs. Nr4a3-Timer Red in CD8^+^ OTI T cells. Splenocytes from Tg4 Nr4a3-Tocky mice were stimulated with 10μM 4Y-MBP, **(C)**, or OTI Nr4a3-Tocky were stimulated with 1μM ova peptide **(D)**, for four hours in the presence of DMSO or 10μM PP2 inhibitor before analysis by flow cytometry for Nr4a3-Timer Blue vs. Nr4a3-Timer Red expression. **(E)** Splenocytes from Tg4 Nr4a3-Tocky mice were stimulated with 10μM 4Y-MBP variant for four hours in the presence of DMSO, 1μM Cyclosporin A (CsA) or 1μM FK506 before analysis of Nr4a3-Time Blue and CD69 expression in CD4+ Tg4 T cells by flow cytometry. Summary data showing the % Nr4a3-Timer Blue^+^ **(F)**, % CD69^+^ **(G)** or CD69 MFI of CD69^+^ **(H)** of CD4^+^ Tg4 T cells, n=5. **(I)** Splenocytes from OTI Nr4a3-Tocky mice were stimulated with 1μM ova peptide for four hours in the presence of DMSO, 1μM Cyclosporin A (CsA) or 1μM FK506 before analysis of Nr4a3-Time Blue and CD69 expression in CD8^+^ OTI T cells by flow cytometry. Summary data showing the % Nr4a3-Timer Blue^+^ **(J)**, % CD69^+^ **(K)** or CD69 MFI of CD69^+^ **(L)** of CD8^+^ OTI T cells, n=3. Statistical analysis by one-way Anova with Tukey’s multiple comparisons test. *p=<0.05, **p=<0.01, ***p=<0.001, ****p=<0.0001. Bars represent mean±SEM.

### Nr4a family members bind NFAT1, but only Nr4a2 and Nr4a3 are sensitive to calcineurin inhibitors

The calcineurin inhibitor data suggested that Nr4a3 is an NFAT responsive distal TCR signalling reporter. Three isoforms of NFAT are expressed in T cells (NFAT1, NFAT2 and NFAT4, (Macian, 2005)). To ascertain NFAT binding, we identified a previously published ChIP-Seq data (GSE64409, (Martinez et al., 2015)) for the binding of NFAT1 to Nr4a family members (Fig. 2A). To control for non-specific binding NFAT1^-/-^ cells were compared to NFAT1 intact mice in unstimulated and stimulated CD8^+^ T cells. Analysis of NFAT1 binding peaks revealed clear NFAT1 binding across the *Nr4a2* and *Nr4a3* genes, which were only evident in stimulated CD8^+^ T cells from NFAT1 wild type mice. *Nr4a1* showed specific peaks upstream of the gene but also displayed considerable non-specific binding throughout the *Nr4a1* gene region. These findings together confirm that Nr4a family members bind NFAT1 within activated CD8^+^ T cells.

**Figure 2:**
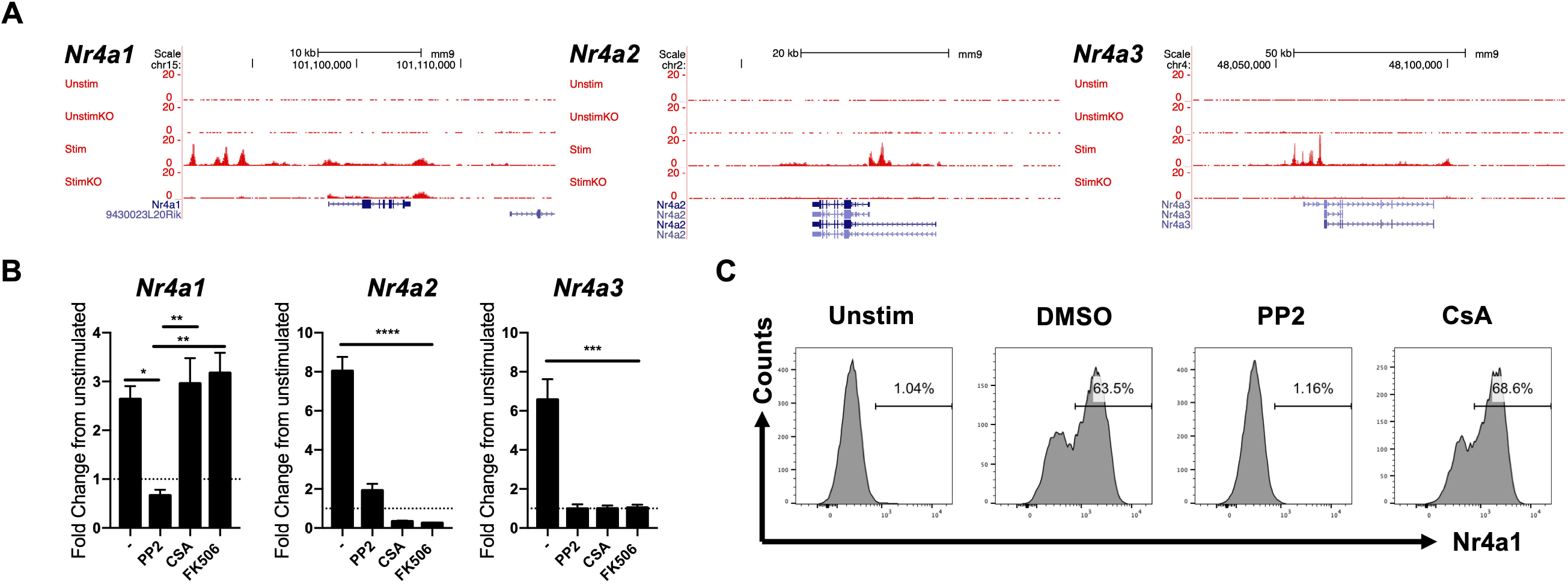
Nr4a family members bind NFAT1, but only Nr4a2 and Nr4a3 are sensitive to calcineurin inhibitors. **(A)** UCSC genome browser tracks showing NFAT1 binding peaks at the *Nr4a1* (left), *Nr4a2* (middle) and *Nr4a3* (right) genes from *in silico* analysis of ChIP-Seq data from GSE64409 in P14^+^ *Tcra*^-/-^ CD8^+^ T cells. Unstim=unstimulated control, UnstimKO= unstimulated NFAT1^-/-^, Stim= PMA and Ionomycin stimulation for 2hrs, StimKO= PMA and Ionomycin stimulation for 1hrs in NFAT1^-/-^ mice. **(B)** Splenocytes from Tg4 Nr4a3-Tocky mice were stimulated with 5μg/ml soluble anti-CD3 for four hours in the presence of DMSO (-), 10μM PP2, 1μM Cyclosporin A (CsA) or 1μM FK506 before extraction of RNA and analysis of fold change in transcription of *Nr4a1* (left), *Nr4a2* (middle) or *Nr4a3* (right) relative to unstimulated condition, n=3. **(C)** Splenocytes from Tg4 Nr4a3-Tocky mice were stimulated with 10uM 4Y-MBP peptide for four hours in the presence of DMSO, 1μM Cyclosporin A (CsA) or 10μM PP2. CD4^+^ T cells were then analysed for intranuclear expression of Nr4a1 protein by flow cytometry. Statistical analysis by one-way Anova with Tukey’s multiple comparison’s test. *p=<0.05, **p=<0.01, ***p=<0.001, ****p=<0.0001. Bars represent mean±SEM.

To test the sensitivity of the other Nr4a family members to calcineurin inhibitors, splenocytes from Tg4 mice were stimulated for four hours with soluble anti-CD3 (to activate CD4^+^ and CD8^+^ T cells) and RNA extracted. Endogenous levels of *Nr4a1, Nr4a2* and *Nr4a3* mRNA were quantified in response to TCR stimulation in order to rule out a potential artefact of the Nr4a3-Tocky BAC transgenic system. Intriguingly, whilst *Nr4a2* and *Nr4a3* showed a complete inhibition of transcriptional upregulation in response to both cyclosporin and FK506 (comparable to the effect of PP2, Src family kinase inhibitor), *Nr4a1* was insensitive to both cyclosporin A and FK506 calcineurin inhibitors (Fig. 2B). Using an antibody to Nr4a1, intracellular staining was performed to verify these findings at the protein level (Fig. 2C). These data suggest that NFAT is necessary for *Nr4a2* and *Nr4a3* expression but is redundant in the regulation of *Nr4a1*.

### ERK signalling is required for optimal Nr4a1, 2 and 3 expression in CD4^+^ and CD8^+^ T cells

NFAT is known to extensively co-operate with AP-1 (Jain et al., 1993; Peterson et al., 1996), and together the two activate many important genes in response to TCR stimulation, such as *Ifng* and *Il2*. AP-1 is dependent on MAP kinase activity (Karin, 1995), in particular ERK pathway activation can lead to the upregulation of c-Fos and enhanced AP-1 transcription factor activity. Past reports have suggested that Nr4a1 is ERK sensitive in ovarian cells (Stocco et al., 2002). To test the sensitivity of *Nr4a3* and CD69 to ERK/AP1 pathway inhibition, splenocytes from Tg4 mice were peptide stimulated in the presence of a MEK/ERK inhibitor (PD0325901, (Barrett et al., 2008)) and compared to the effects of DMSO or cyclosporin A (Fig. 3A). TCR induction of Nr4a3 was, as previously shown, abolished by CsA, but also showed a significant attenuation in the presence of ERK pathway inhibition (Fig. 3A&B). CD69 levels were partially inhibited by CsA but showed a profound reduction in the presence of ERK pathway inhibitors (Fig. 3C). Stimulation of CD4^+^ Tg4 T cells in the presence of both inhibitors resulted in complete inhibition of CD69 expression (Fig. 3C&D).

**Figure 3:**
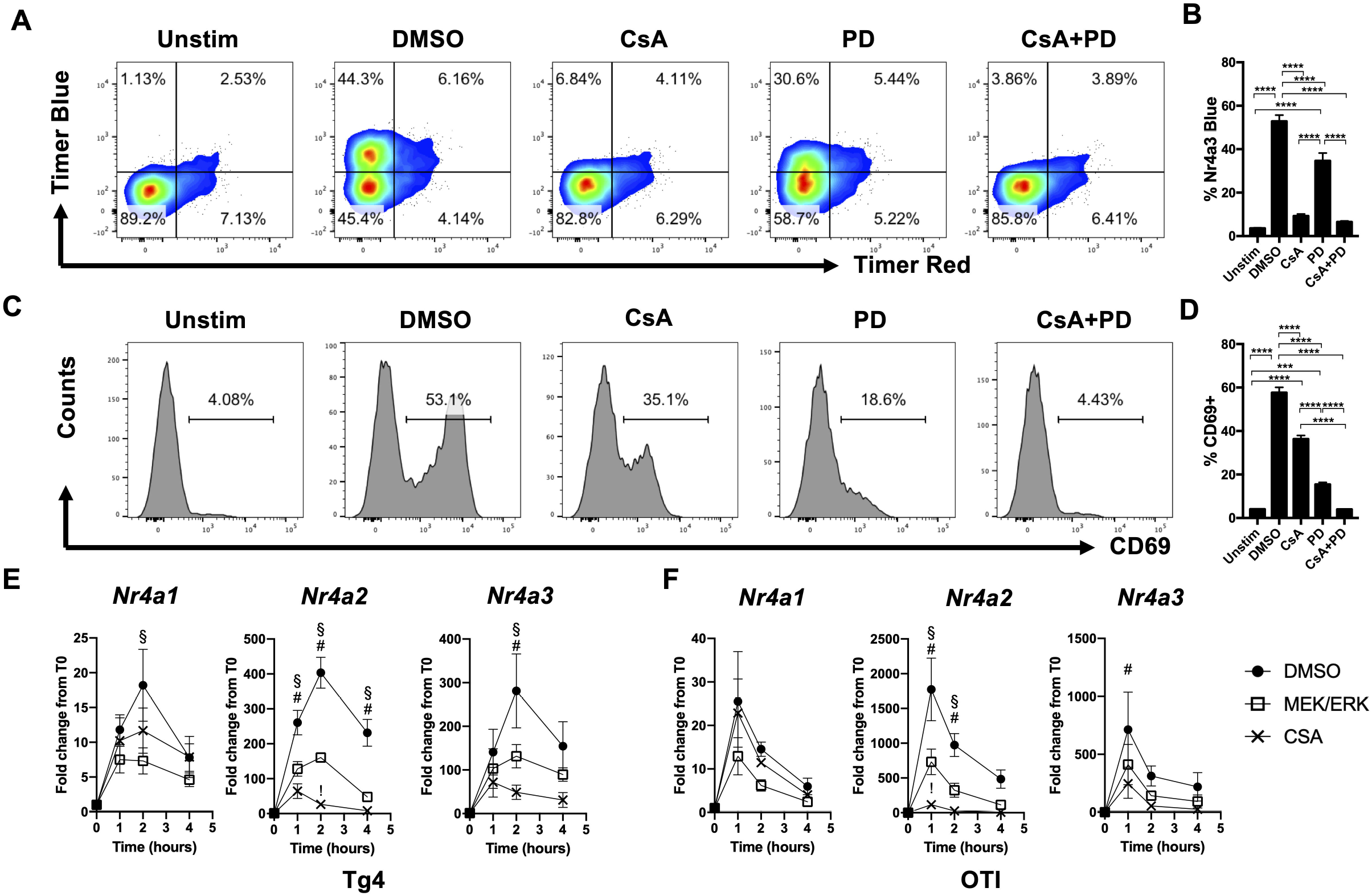
ERK signalling is required for optimal *Nr4a1, 2 and 3* expression in CD4^+^ and CD8^+^ T cells. **(A)** Splenocytes from Tg4 Nr4a3-Tocky mice were stimulated with 10μM 4Y-MBP variant for four hours in the presence of DMSO, 1μM Cyclosporin A (CsA, NFAT pathway inhibitor), 5μM PD0325901 (PD) MEK inhibitor or 1μM CsA and 5μM PD before analysis of Nr4a3-Time Blue vs. Nr4a3-Timer Red. **(B)** Summary data of data from **(A)**, n=5-6, statistical analysis by one-way Anova with Tukey’s multiple comparisons test. **(C)** histogram expression of CD69 in CD4^+^ Tg4 T cells stimulated with 10μM 4Y-MBP variant for four hours in the presence of DMSO, 1μM Cyclosporin A (CsA), 5μM PD0325901 (PD) MEK inhibitor or CsA +PD by flow cytometry. **(D)** Summary data from (C), n=5, statistical analysis by one-way Anova with Tukey’s multiple comparisons test. Splenocytes from Tg4 Nr4a3-Tocky mice were stimulated with 10μM 4Y-MBP variant **(E)** or OTI Nr4a3-Tocky mice with 1μM ova peptide **(F)** for four hours in the presence of DMSO, 1μM Cyclosporin A (CsA) or 5μM PD0325901 (PD) and RNA extracted at 0, 1, 2 and 4hrs following culture. Graphs depict fold change in Nr4a1, Nr4a2 and Nr4a3 mRNA levels relative to T=0hrs in DMSO treated (black circles), CsA treated (Black crosses) or PD0325901 treated (open squares), n=4 (E) or n=3 (F). Bars represent mean±SEM. #=CsA treated significantly reduced from DMSO; §=PD treated significantly reduced from DMSO; !=CsA treated significantly reduced from PD. Statistical analysis by two-way Anova with Tukey’s multiple comparison’s test.

To further analyse the kinetics of *Nr4a* gene family transcription in response to TCR stimulation, time course analysis was performed on transcript levels in peptide stimulated Tg4 CD4^+^ and OTI CD8^+^ T cells (Fig. 3E&F) in the presence of NFAT or ERK pathway inhibitors. These findings revealed significant inhibition of all *Nr4a* family members by ERK pathway inhibition at two hours post stimulation, but no significant effect of NFAT pathway inhibition on *Nr4a1* at any time point. In contrast, NFAT pathway inhibition resulted in a greater repression of *Nr4a2* and *Nr4a3* transcription compared to the ERK pathway (Fig. 3E&F), in keeping with the analysis of Nr4a3-Timer expression by flow cytometry (Fig. 3A). OTI CD8^+^ T cells largely mirrored Tg4 CD4^+^ T cells in the sensitivity of Nr4a members to NFAT and ERK pathway inhibitors, with all showing partial ERK dependence, but only *Nr4a2* and *Nr4a3* being sensitive to calcineurin inhibitors (Fig. 3E). These findings were verified by flow cytometric analysis in CD8^+^ OTI Nr4a3-Tocky mice, revealing that OTI CD8^+^ T cells also require ERK pathway signalling for optimal Nr4a3 expression, but are absolutely dependent on the CaN/NFAT pathway (Supplementary Fig. 2).

### Constitutively active NFAT1 is sufficient to induce *Nr4a2* and *Nr4a3* expression

The previous findings suggested that Nr4a2 and Nr4a3 are NFAT-dependent transcription factors, which require ERK signalling for optimal expression. Nr4a1 on the other hand was not dependent on the NFAT pathway, but had a partial dependency on ERK activity. Given that NFAT:AP-1 complexes are critical to many facets of T cell biology, we were interested to know whether NFAT activity alone would be sufficient to drive *Nr4a2* and *Nr4a3* expression. We therefore performed *in silico* analysis of a previously published RNA-seq dataset of T cells expressing a constitutively active form of NFAT1 that was modified to abrogate its binding to AP1 ((Martinez et al., 2015) CA-RIT-NFAT1, Fig. 4). This system permits an analysis of genes that can be activated solely by the presence of activated NFAT1. Analysis of *Nr4a* family members revealed that CA-RIT-NFAT1 resulted in a statistically significant upregulation of Nr4a3 within CD4 (>13-fold, Fig. 4A) and CD8 T cells (>50-fold, Fig. 4B) and Nr4a2 within CD8s (>15-fold, Fig. 4B). Therefore, NFAT1 activity alone is sufficient for the induction of *Nr4a2* and *Nr4a3* expression, but not *Nr4a1* and *Cd69*, in response to TCR signalling.

**Figure 4:**
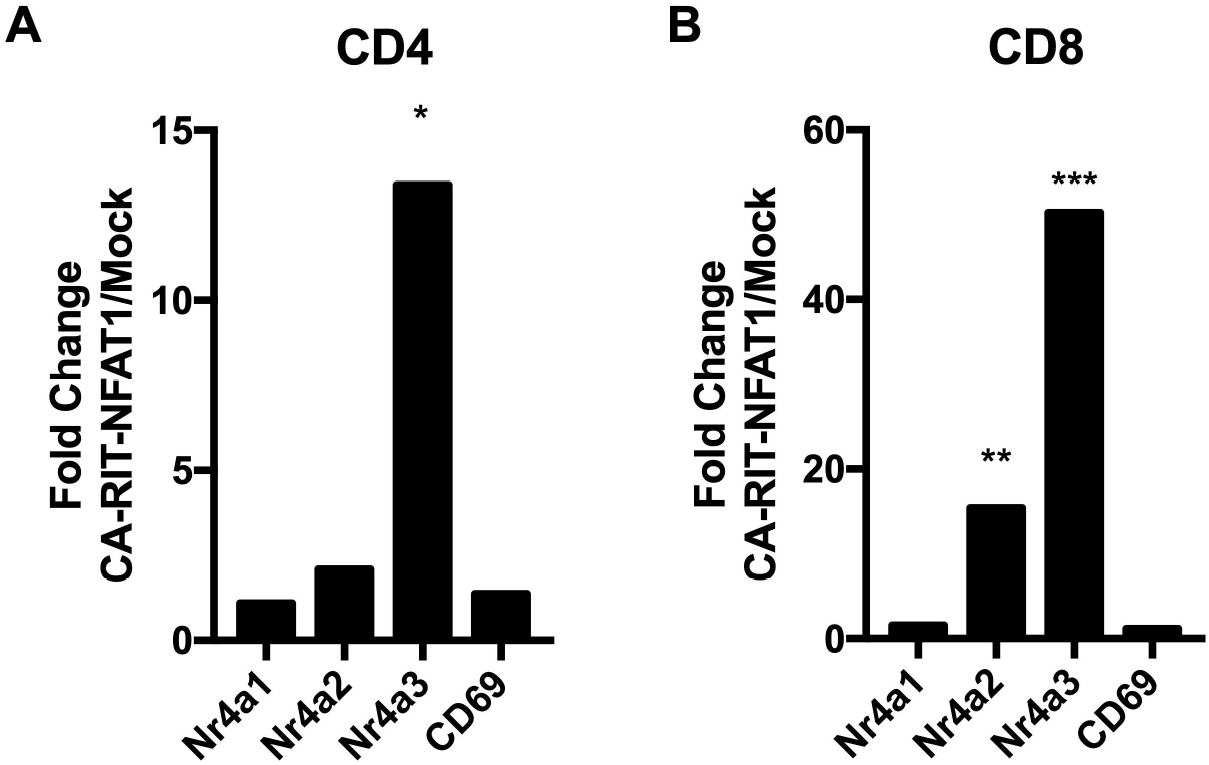
Constitutively active NFAT1 is sufficient to induce *Nr4a2* and *Nr4a3* expression. Analysis of RNA-seq data from GSE64409 displaying the fold change in expression of *Nr4a1, Nr4a2, Nr4a3* and *Cd69* in **(A)** CD4^+^ or **(B)** CD8^+^ T cells transduced with a constitutively active form of NFAT1 that is unable to interact with AP-1 (CA-RIT-NFAT1) vector compared to mock vector. Adjusted p values based on DESeq analysis * = p=0.0206, ** = p=0.00305, *** = p=0.00016.

### Nr4a1 and Nr4a3 expression patterns distinguish tonic versus activated lymphocyte signalling events in vivo

Given the differential sensitivity of Nr4a1 and Nr4a3 to TCR signalling pathways, we interrogated their expression in vivo by crossing Nur77GFP to Nr4a3-Tocky mice. Using Nur77-GFP, we confirmed that the Nur77GFP BAC transgene was also insensitive to calcineurin inhibitors, but did display a partial regulation through ERK in CD4^+^ and CD8^+^ T cells (Fig. 5A). T cell development in the thymus was analysed to understand the expression patterns of Nr4a1 and Nr4a3 that occur during TCR-mediated selection events (Fig. 5B). Nur77GFP was largely absent within pre- selection immature TCRβ^lo^ double positive (DP) cells, but increased in expression in the intermediate TCRβ^hi^ DP fraction indicative of tonic signalling following the initiation of positive selection, such that all positively selected CD4 single positive (SP) cells expressed Nur77GFP. Nr4a3 expression was largely absent from the DP and CD4^+^CD25^-^ fraction, however it was expressed within the CD4SP CD25^hi^ fraction, which is enriched with Treg and Treg precursor cells. Comparison of Nur77GFP and Nr4a3-Timer Red revealed how Nr4a3-Timer expression only emerged at the highest levels of GFP, highlighting that the threshold for activation in vivo is higher for Nr4a3 than Nr4a1. Importantly, and in contrast to tonic in vivo TCR signalling, administration of cognate peptide in Tg4 Nr4a3-Tocky mice elicited Nr4a3 expression in both DP and CD4SP thymocytes. This suggests that the lack of Nr4a3 expression in DP and CD4SP thymocytes is not due to inaccessibility of the Nr4a3 locus (Supplementary Fig 3). Rather, unlike Nur77, Nr4a3 expression requires a higher TCR signalling strength which is achieved through cognate peptide:MHC mediated T cell activation. In keeping with the thymus, peripheral CD4^+^CD25^-^ T cells expressed intermediate Nur77GFP levels and largely lacked Nr4a3 expression. In contrast CD4^+^CD25^+^ T cells were enriched for Nr4a3^+^ T-cells as previously reported (Bending et al., 2018b)(Fig. 5C).

**Figure 5:**
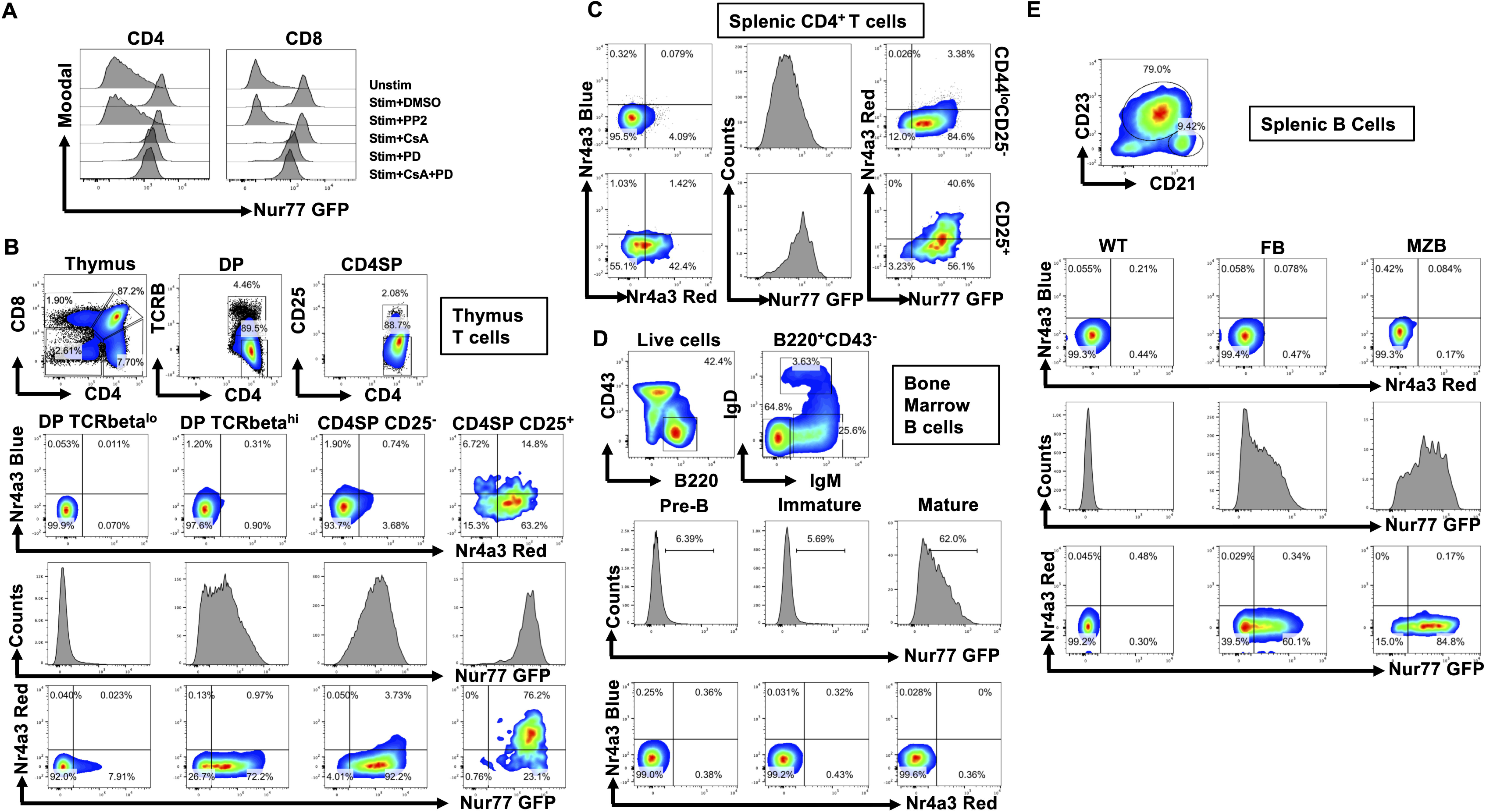
Nr4a1 and Nr4a3 expression patterns distinguish tonic versus activated lymphocyte signalling events in vivo. **(A)** Splenocytes from Nur77-GFP mice were stimulated with 3μg/ml anti-CD3 for four hours in the presence of DMSO, 1μM Cyclosporin A (CsA), 5μM PD0325901 (PD) MEK inhibitor or 1μM CsA and 5μM PD before analysis of Nur77GFP levels in CD4^+^ or CD8^+^ T cells by flow cytometry. **(B)** Thymus from Nur77-GFP Nr4a3-Tocky mice were analysed for expression of Nur77GFP and Nr4a3-Timer blue and Nr4a3-Timer Red expression within TCRβ^lo^ and TCRβ^hi^ DP and CD25^-^ and CD25^+^ CD4SP subsets by flow cytometry. **(C)** Splenic CD4^+^CD25^-^ and CD4^+^CD25^-^ T cells were analysed from Nur77-GFP Nr4a3-Tocky mice for expression of Nur77-GFP and Nr4a3-Timer Blue and Nr4a3-Timer Red expression by flow cytometry. **(D)** Bone marrow from Nur77-GFP Nr4a3-Tocky mice was analysed for expression of Nur77GFP and Nr4a3-Timer blue and Nr4a3-Timer Red expression within B220^+^CD43^-^ B cells. Gating on pre-B cells (B220^+^CD43^-^IgM^-^IgD^-^), immature (B220^+^CD43^-^IgM^+^IgD^-^) and mature B cells (B220^+^CD43^-^IgM^+^IgD^+^) **(E)** Splenic B cell from Nur77-GFP Nr4a3-Tocky mice (gated on B220^+^CD19^+)^ were divided into follicular (FB, CD21^+^CD23^-^) and marginal zone (MZB, CD21^+^CD23^+^) subsets and analysed for expression of Nur77-GFP and Nr4a3-Timer Blue and Nr4a3-Timer Red expression by flow cytometry.

Nur77GFP is also known to be expressed in developing and mature B cells (Moran et al., 2011; Zikherman et al., 2012). Similarly, we found that mature B cells expressed Nr4a1, but not Nr4a3, highlighting that tonic BCR signalling can also activate Nr4a1 but not Nr4a3 expression (Fig. 5D), and in the spleen, marginal zone (MZ) B cells exhibited higher levels of Nur77GFP than follicular (F) B cells, whilst continuing to be absent for Nr4a3-Timer expression (Fig. 5E). These findings highlight that, as with T cells, tonic BCR signalling is able to elicit Nr4a1 but not Nr4a3 expression, whilst co-expression requires crosslinking of the BCR (Bending et al., 2018b).

## Discussion

The data have highlighted how the differential sensitivities of Nr4a1 and Nr4a3 to distal TCR signalling pathways can be exploited to identify T cells undergoing different types of physiological antigen receptor signalling in vivo. Nr4a1 exhibits a lower threshold for activation, which is triggered within DP T cells upon beta selection in the thymus and upon co-expression of IgM on B cells in the bone marrow. Nr4a3 requires full activation of the calcineurin/NFAT pathway, which occurs during thymic Treg development but also by cognate peptide-MHC interaction in vivo (Supplementary Fig. 4). Nr4a1 appears to be expressed upon successful rearrangement of antigen receptors during lymphocyte development.

Nr4a receptors are emerging as an exciting target for immunomodulation (Flemming, 2019), in particular as a potential strategy to fine-tune CAR T cell attack of solid tumours (Li and Zhang, 2019). Here we have shown that *Nr4a1, Nr4a2* and *Nr4a3* are distinctly regulated by the CaN/NFAT pathway, but that all three transcription factors require MEK/ERK pathway activation for optimal expression in response to TCR stimulation in both CD4^+^ and CD8^+^ T cells. Interestingly, analysis of NR4A1 protein regulation in anti-CD3 stimulated human T cells has shown that they are also insensitive to calcineurin inhibitors (Ashouri and Weiss, 2017), but show a partial ERK dependence, suggesting that regulation of Nr4a1 may be conserved between mice and humans.

Remarkably, despite being insensitive to calcineurin inhibitors, the *Nr4a1* gene region exhibited binding of NFAT1. Surprisingly, both cyclosporin and FK506 had no effect on the transcriptional dynamics of *Nr4a1* expression in response to peptide stimulation of either CD4^+^ and CD8^+^ T cells. This could represent redundancy in the regulation of Nr4a1 by NFAT1, given that Nr4a1 appears more sensitive to a broad range of distal TCR signalling pathways. In particular, it has been previously observed that cyclosporin may interfere with Nr4a1 biology at the level of its DNA binding activity through its N-terminal protein region (Yazdanbakhsh et al., 1995), without interfering with its transcription. In this way, cyclosporin may abrogate the biological effects of all Nr4a family members through disparate mechanisms of action. On the other hand, constitutively active NFAT1 did not significantly alter *Nr4a1* expression in CD4^+^ or CD8^+^ T cells transduced with the CA-RIT-NFAT1 vector, suggesting that *Nr4a1* cannot be activated by NFAT1 activity alone. However, as the CA-RIT-NFAT1 construct is incapable of binding AP-1 (Martinez et al., 2015), it is still possible that NFAT:AP1 complexes could redundantly activate Nr4a1, or that NFAT2, and NFAT4 may play roles. For instance, *Cd69* was not significantly upregulated in CA-RIT-NFAT1 T cells but is abolished by coadministration of CaN inhibitors and MEK/ERK pathway inhibitors. These suggest that CD69, like Nr4a1, has multiple redundant mechanisms controlling its transcription in T cells downstream of TCR signalling, including a partial regulation by JAK inhibitors (Ashouri and Weiss, 2017).

The discovery that ERK has a role in optimal Nr4a expression is in keeping with studies in other tissues. NOR1 (Nr4a3) has been shown to be regulated by the MAPK/ERK pathway in response to platelet derived growth factor in vascular smooth muscle cells (Nomiyama et al., 2006). Similarly, in ovarian cells, calcium-dependent activation of ERK mediates AP-1 induction of Nur77 (Nr4a1) (Stocco et al., 2002), again establishing a common link between ERK signalling and the regulation of Nr4a expression in a wide range of cell types.

The finding that all Nr4a members are regulated by the Src family kinase suggests that the initiation of the pathway to Nr4a induction is by the activity of lymphocyte-specific protein tyrosine kinase (Lck). Lck associates with the cytoplasmic tails of the CD4 and CD8 co-receptors (Veillette et al., 1988). Peptide-MHC engagement of the TCR leads to Lck mediated phosphorylation of the TCR and CD3 chains (Straus and Weiss, 1992). Interestingly, past work has suggested that more CD4 coreceptors are loaded compared to CD8 molecules within T cells, which can affect the dwell time of T cells in response to negatively selecting antigens (Stepanek et al., 2014). Comparison of *Nr4a* transcriptional dynamics in this study suggested that CD8^+^ T cells receive an initial shorter, sharper and stronger activation of *Nr4a* transcription, which peaks at 1hr. Interestingly *Nr4a* receptor transcription in CD4^+^ T cell peaked around 2hrs and appeared to plateau. These findings warrant further investigation to understand the temporal regulation of Nr4a expression following antigen experience, but could also reflect differences in early negative feedback pathways, differential affinity of the TCR and antigen, and/or differences in MHC Class I and II expression levels.

Despite the different sensitivities of Nr4a1 and Nr4a2/3 to NFAT pathway inhibitors, the overall kinetics of expression in peptide stimulated CD4^+^ or CD8^+^ T cells was fairly uniform. However, it is important to note that these readouts are from primary stimulation of a largely naïve population of TCR transgenic T cells. For instance, antigen-experienced T cells in humans show altered ERK and p38 phosphorylation patterns compared to naïve T cells (Adachi and Davis, 2011). Nr4a family receptors may therefore be useful in studying the re-tuning of T cell receptor signalling pathways – in particular comparing Nr4a1 to Nr4a2/Nr4a3 transcriptional responses. Currently within the field there exist several Nr4a1 (Nur77) GFP transgenic reporters (Moran et al., 2011; Zikherman et al., 2012). These reporters have been used to study Treg and iNKT development, as well as address B cell activation to antigen. However, as previously alluded to by Weiss and colleagues, temporal analysis of Nr4a1-GFP reporters may be hampered by the persistence of GFP expression following antigen encounter (Au-Yeung et al., 2014), and its induction by tonic signalling events during development. The Nr4a3-Tocky system employed in this study does not suffer from this same issue, since the half-life of Timer Blue protein is 4hrs (Bending et al., 2018b), allowing a sensitive readout for activating TCR signalling over much shorter periods of time. Given that NFAT in the absence of AP-1 induces a chronic T cell exhaustion phenotype, such as PD1^high^ Lag3^high^ CD8^+^ T cells (Martinez et al., 2015), it is likely that Nr4a3-Tocky will prove a useful model for understanding NFAT pathway activity in animal models of T cell dysfunction and cancer in vivo. Furthermore, assessing Nr4a3 and Nr4a1 co-regulation will be a potent tool to interrogate alterations in TCR signalling in T cells during their development or as a T cell intrinsic readout for the effects of immunotherapies.

## Acknowledgements

Work funded by the Wellcome Trust Seed award (D.B., E.J., T.A.E.E). S.K. was supported by an Arthur Thomson summer studentship. M.O. was supported by a BBSRC David Phillips Fellowship. D.W. is supported by the University of Birmingham. D.B. is supported by a University of Birmingham Fellowship. K.M.T. and J.Y.P. were supported by the BBSRC (BB/S003800/1). GA is supported by an MRC Programme Grant (MR/N000919/1).

## Author Contributions

D.B. designed and conceptualised the experiments, performed in silico analyses and wrote the paper. E.K.J, T.A.E.E, N.T. and S.K. conducted experiments. D.C.W provided Tg4 Tiger mouse line and M.O. provided Nr4a3-Tocky strain. G.A. provided Nur77-GFP mouse line and expert advice on thymic T cell development. J.Y.P and K.M.T. provided reagents and expert advice on flow cytometric staining for B cell development.

## Declaration of Interests

The authors declare no competing interests.

## Methods

### Mice

Nr4a3-Tocky Tg4 Tiger mice were used as the F1 generation by crossing Nr4a3-Tocky ((Bending et al., 2018b) founder line 323, obtained from Dr. Masahiro Ono, Imperial College London under MTA) mice with the Tg4 TCR IL-10-GFP Tiger (Burton et al., 2014; Kamanaka et al., 2006) provided by Prof. David Wraith, UoB. Nr4a3-Tocky Great-Smart17A (Price et al., 2012) were initially bred to homozygous OT1 mice (purchased from Charles River laboratories) to generate F1 OT1 Nr4a3-Tocky Great Smart-17A mice. Nur77GFP mice (Moran et al., 2011) were used alone or bred to Nr4a3-Tocky mice to generate Nur77GFP Nr4a3-Tocky. All animal experiments were performed in accordance with local Animal Welfare and Ethical Review Body at the University of Birmingham and under the authority of a Home Office project licence, P18A892E0A.

### In *vitro* cultures

Single cell suspensions of splenocytes from Nr4a3-Tocky Tg4 Tiger and OTI Nr4a3-Tocky mice were generated by forcing organs through 70-μm cell strainers (Corning). For spleens, a red blood cell (RBC) lysis stage was performed (Invitrogen) according to manufacturer’s instructions. Cells were washed once and cultured at 1 x 10^6^ cells per well on 96-well U-bottom plates (Corning) with or without the presence of peptides or anti-CD3 and inhibitors in a final volume of 200 μl RPMI1640 + L-glutamine (Gibco) containing 10 % FCS and penicillin/streptomycin (Life Technologies). For Tg4 stimulation 10μM of MBP Ac1-9[4Y] peptide (which is a modified form of the acetylated native N-terminal murine MBP peptide (Ac-ASQYRPSQR, GL Biochem LTD Shanghai, China), for OTI stimulation 1μM ova peptide (SIINFEKL, Cambridge BioScience) or soluble anti-CD3 (145-2C11, BioLegend). Cells were incubated at 37°C and 5% CO_2_ and analysed at the indicated time points for RNA expression or flow cytometric analysis.

### Inhibitors

The following inhibitors were dissolved in DMSO (Sigma) and used: Cyclosporin A (Cambridge Bioscience, 1μM), FK506 (Cayman Chemical Company, 1μM), PD0325901 (Cambridge Bioscience, 5μM), PP2 (Sigma, 10μM) and DMSO (Sigma, 0.1%).

### Flow cytometric analysis

For analysis of thymus and splenic lymphocytes single cell suspensions were prepared as described above. For analysis of bone marrow B cells, femurs were flushed with media and then RBC lysis performed before staining for flow cytometric analysis. Cells were washed once and stained in 96-well U-bottom plates (Corning). Analysis was performed on a BD LSR Fortessa X-20 instrument. The blue form of the Timer protein was detected in the blue (450/40 nm) channel excited off the 405 nm laser. The red form of the Timer protein was detected in the mCherry (610/20) channel excited off the 561 nm laser. A fixable eFluor 780-flurescent viability dye (eBioscience) was used for all experiments. The following directly conjugated antibodies were used in these experiments: CD4 Alexa Fluor 700 (Clone RM4-4, BioLegend), CD4 BUV737 (Clone GK1.5, BD Biosciences) TCR Vβ8.1, 8.2 PerCP-eFluor 710 (Clone KJ16-133, Thermofisher), TCR Vβ8.1/8.2 BUV395 (Clone MR5-2, BD Biosciences), TCRβ AF700 (Clone H57-597, BioLegend), CD69 APC (Clone H1.2F3, BioLegend), CD8 Alexa Fluor 700 (Clone 53-6.7, BioLegend), CD8 BUV395 (Clone. 53-6.7, BD Biosciences), TCR Vα2 PerCP/Cyanine5.5 (Clone B20.1, BioLegend), TCRVβ5.1/5.2 APC (Clone MR9-4, BioLegend), B220 PerCPcy5.5 (Clone RA3-6B2, BD Biosciences), CD43 biotin (Clone S7 BD Biosciences), streptavidin BV711 (BioLegend), IgM PE-Cy7 (Clone RMM-1, BioLegend), IgD APC (Clone 11-26C, Invitrogen), CD21 APC (Clone 7G6, BD Biosciences), CD23 PE-Cy7 (Clone B3B4, eBioscience), CD25 PerCPCy5.5 (PC61, BioLegend), CD19 BUV737 (Clone 1D3, BD Biosciences). For staining of Nr4a1 protein, cells were fixed and stained with Nr4a1 PerCP-eFluor 710 (Clone 12.14, eBioscience) using the eBioscience Foxp3/Transcription factor staining buffer set according to the manufacturer’s instructions (ThermoFisher Scientific).

### In vivo peptide immunisation

Tg4 Nr4a3-Tocky mice were immunised with 80μg of MBP Ac1-9[4Y] sub cutaneous or vehicle control (PBS). 4hrs later, thymus was removed and analysed for expression of Nr4a3-Timer Blue and Red.

### qPCR analysis

Following *in vitro* cultures, RNA was extracted using the RNeasy kit (QIAGEN) according to the manufacturer’s instructions. cDNA was generated using random hexamers and Superscript IV (Invitrogen) according to the manufacturer’s instructions. mRNA expression was quantified using PowerUp SYBR green (Life Technologies) and normalised to the housekeeping gene *Hprt* using the 7900HT sequence detection system or 7500 (Applied Biosystems). Fold change in expression was calculated using the delta-deltaCt method. Primer sequences: *Hprt* For: AGCCTAAGATGAGCGCAAGT, *Hprt* Rev: TTACTAGGCAGATGGCCACA, *Nr4a1* For: TGTGAGGGCTGCAAGGGCTTC, *Nr4a1* Rev: AAGCGGCAGAACTGGCAGCGG (Sekiya et al., 2013), *Nr4a2* For: CTGTGCGCTGTTTGCGGTGAC, *Nr4a2* Rev: CGGCGCTTGTCCACTGGGCAG (Sekiya et al., 2013), *Nr4a3* For: AGGGCTTCTTCAAGAGAACGG, *Nr4a3* Rev: CCATCCCGACACTGAGACAC (designed using NCBI Primer Blast).

### *In silico* ChIP-Seq

Processed bigwig data files deposited in GSE64409 (Martinez et al., 2015) were downloaded and hosted in CyVerse Discovery Environment (de.cyverse.org/de) and then mapped against the mm9 genome using the UCSC genome browser. These files contain analysis of NFAT1 ChIP-Seq in CD8 T cells from WT and NFAT1^-/-^ mice either unstimulated or stimulated for 1hr with PMA and Ionomycin as described in (Martinez et al., 2015).

### RNAseq analysis

Log2 fold change estimates of *Nr4a1, Nr4a2, Nr4a3* and *Cd69* expression was extracted from DESeq data deposited in (Martinez et al., 2015) for CD4^+^ or CD8^+^ T cells either transfected with mock vector or CA-RIT-NFAT1.

### Statistical analysis and data visualisation

Statistical analysis was performed on Prism 7 or 8 software (GraphPad). For comparison of more than two means a one-way ANOVA with Tukey’s multiple comparisons test was used. For comparison of more than two means over time, a two-way ANOVA with Tukey’s multiple comparison’s test was used. Adjusted p values for Figure 5 were extracted from DESeq analysis in (Martinez et al., 2015). Variance is reported as mean±SEM unless otherwise stated. *p=<0.05, **p=<0.01, ***p=<0.001, ****p=<0.0001.

## Supplementary Figure Legends

**Supplementary Figure 1: Analysis of Tg4 Nr4a3-Tocky and OTI Nr4a3-Tocky mice**

**(A)** Nr4a3-Tocky mice were crossed with Tg4 Tiger mice to generate Tg4 Tiger Nr4a3-Tocky mice, which are specific for myelin basic protein-derived peptides. Thymus (top) and spleen (bottom) were analysed for CD4 vs CD8 expression within live lymphocytes (left), CD4 vs. TCRVβ8.1/8.2 (middle) gated on CD4SP cells, and Nr4a3-Timer Blue vs Nr4a3-Timer Red in CD4SPVβ8.1/8.2^+^ T cells. **(B)** Nr4a3-Tocky Great Smart-17A mice were bred with OTI TCR transgenic mice, which are specific for ova peptide. Thymus (top) and spleen (bottom) were analysed for CD4 vs CD8 expression within live lymphocytes (left), TCRVβ5.1/5.2 vs TCRVα2 (middle) gated on CD8SP cells, and Nr4a3-Timer Blue vs Nr4a3-Timer Red in CD8SPVβ5.1/5.2^+^Vα2^+^ T cells.

**Supplementary Figure 2: CD8^+^ T cells require ERK signalling for optimal Nr4a3-Timer expression**

**(A)** Splenocytes from OTI Nr4a3-Tocky mice were stimulated with 1μM ova peptide for four hours in the presence of DMSO, 1μM Cyclosporin A (CsA), 5μM PD0325901 (PD) MEK inhibitor or 1μM Cyclosporin A (CsA)+5μM PD0325901 (PD) MEK inhibitor before analysis of Nr4a3-Timer Blue vs. Nr4a3-Timer Red expression within CD8^+^ OTI T-cells. **(B)** Summary data from (A), n=4, bars represent mean±SEM. Statistical analysis by one-way Anova with Tukey’s multiple comparisons test.

**Supplementary Figure 3: Peptide administration activated Nr4a3 expression in developing thymocytes**

Tg4 Nr4a3-Tocky mice were immunised with 80ug of MBP [4Y] peptide or vehicle control (PBS) and thymus removed 4hrs later and TCRβ^lo^ and TCRβ^hi^ DP and CD4SP subsets analysed for the expression of Nr4a3-Timer Blue vs Timer Red by flow cytometry.

**Supplementary Figure 4: Differential regulation of Nr4a members by ERK and NFAT signalling distinguishes tonic from activating TCR signalling events in vivo.**

Nr4a family members depend on Src family kinase activity (Lck), which leads to the downstream activation of MEK/ERK and calcineurin (CaN) and NFAT pathways. Nr4a2 and Nr4a3 require CaN activity and NFAT1 binds Nr4a2 and Nr4a3 and is sufficient to promote their transcription. All Nr4a family members are sensitive to the ERK pathway, most likely through AP-1 transcription factor activity (dashed lines imply partial dependence, bold lines complete dependence on pathway). Nr4a1 shows enhanced sensitivity to TCR signalling suggesting multiple redundant pathways lead to its activation (suggested by question mark). Nr4a1 can be induced by tonic signalling alone, whilst Nr4a3 requires a higher threshold of signalling through activating TCR signalling.

