## Supplementary Figures 1-4 for "Differential Nr4a1 and Nr4a3 expression discriminates tonic from activated TCR signalling events in vivo"

### Fig S1: Analysis of Tg4 Tiger Nr4a3-Tocky and OTI Nr4a3-Tocky mice

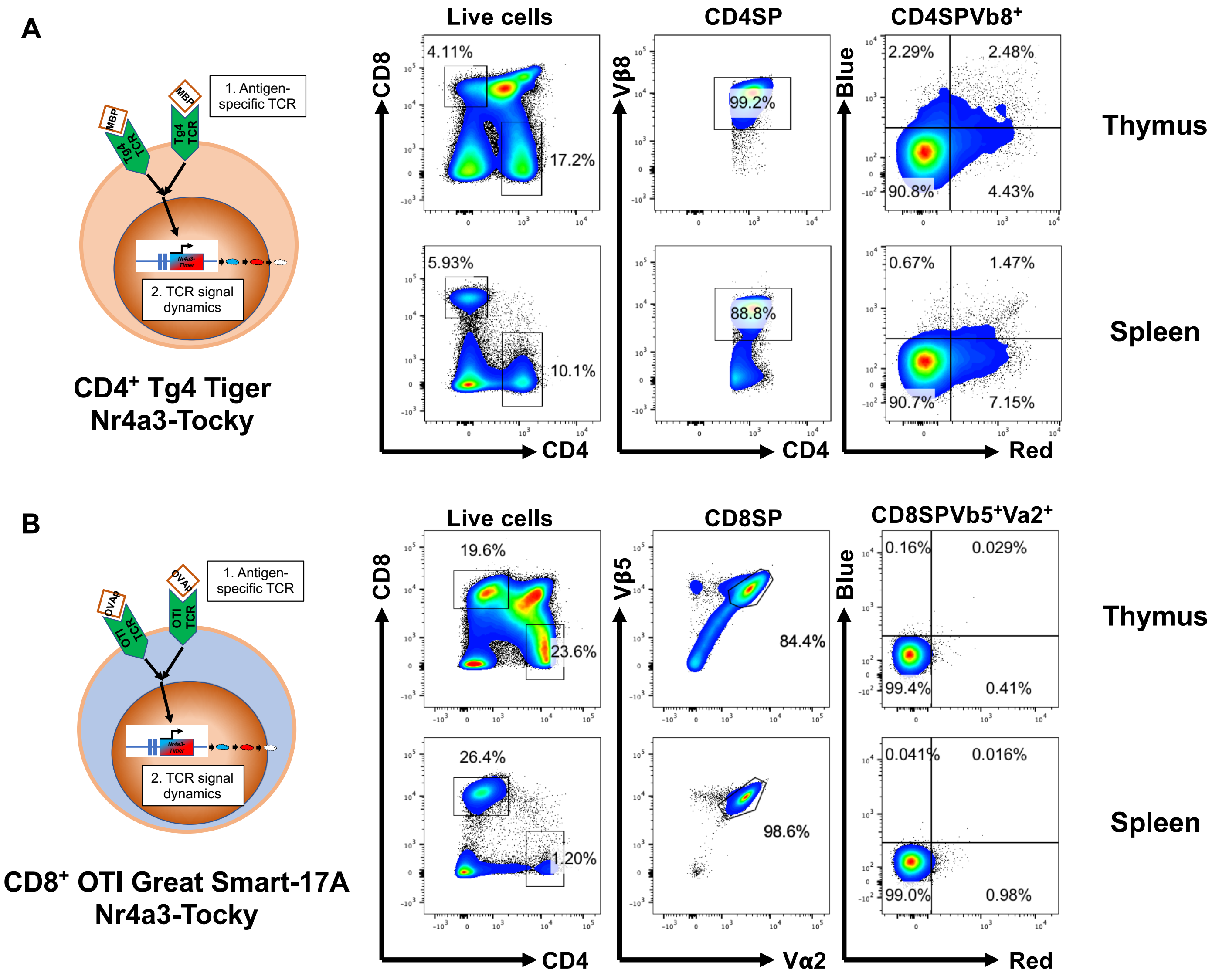

### Fig S2: CD8⁺ T cells require ERK signaling for optimal Nr4a3 expression

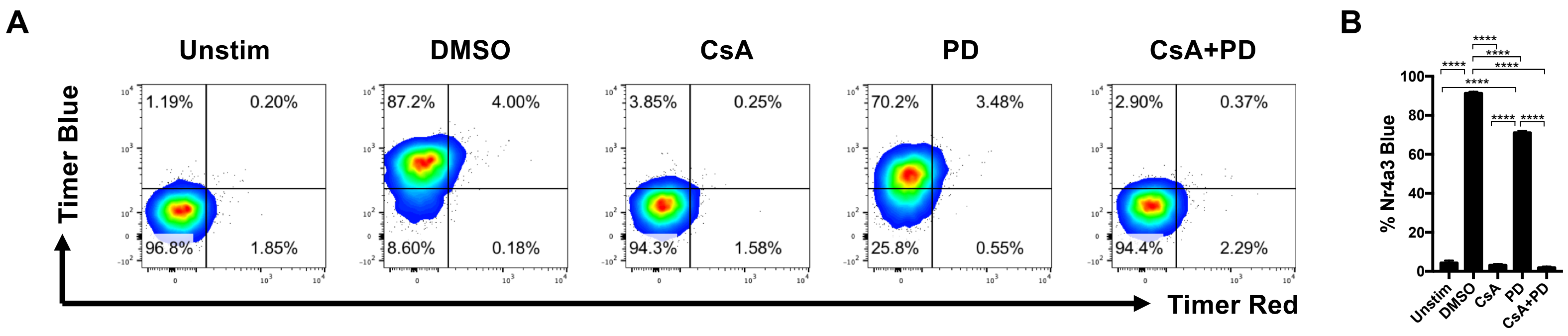

**Fig S3: Peptide administration activates Nr4a3 expression in developing thymocytes**

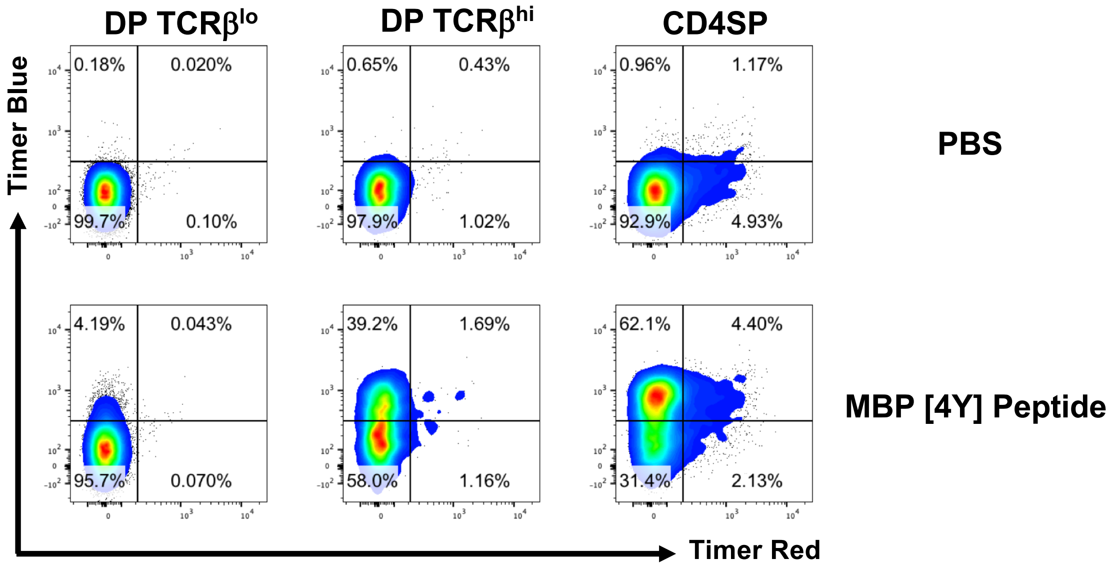

Fig S4: Graphical abstract

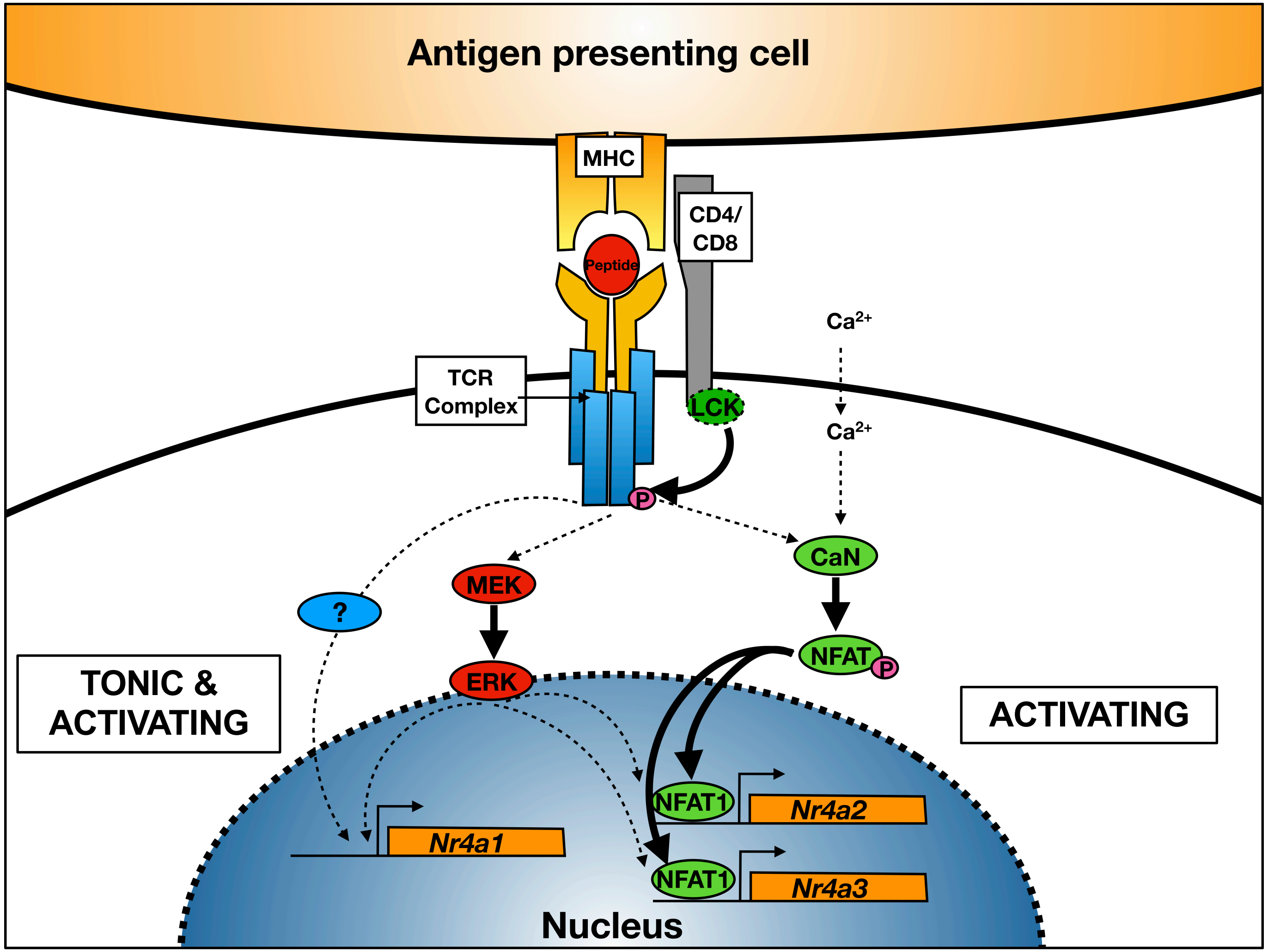
